## Supplemental materials for "Self-Supervised Learning Reveals Clinically Relevant Histomorphological Patterns for Therapeutic Strategies in Colon Cancer"

### Table 1: Tissue annotation for 47 HPCs

| HPC | Super-cluster | Label | General description | Tumor epithelium | Tumor stroma | Immune cells | Other | HPC tissue type composition |
|---|---|---|---|---|---|---|---|---|
| 0 | Tumor stroma | Disorganized stroma | Mostly disorganized stroma, sometimes some aligned strands and sometimes some muscle tissue, immune cells in between, with a few isolated tumor cells | A couple isolated tumor cells | Primarily tumor stroma on tiles. Stroma-high tiles, more disorganized than aligned stroma | Some immune cells in hotspots, immune-high overall | Some muscle tissue | Stroma 70%, Muscle tissue 20%, Immune cells 10% |
| 1 | Tumor stroma | Edematous stroma | Loose, edematous and disorganized stroma with background, sometimes vessels and an artefact. Most likely not tumor-induced stroma but healthy serosa as there is also some fatty tissue | A couple isolated tumor cells | Primarily loose, edematous and disorganized stroma-high tiles with background, sometimes vessels with some erythrocytes. Most likely not tumor-induced stroma but healthy serosa looking at pattern, and there is also some fatty tissue | A couple immune cells, immune-low | Some fatty tissue | Stroma 80%, Fatty tissue 10%, Background 10% |
| 2 | Tumor stroma | Aligned stroma - well differentiated tumor | Mostly up until 50% stroma in aligned, straight strands. Well differentiated tumor in between, with a little dirty necrosis | Mostly well differentiated tumor epithelial cells | Mostly aligned, straight strands of tumor stroma on the stroma-low tiles | A couple immune cells, and some (dirty) necrosis, immune-low | N/A | Stroma 40%, Tumor epithelial cells 60% |
| 3 | Tumor epithelium | Moderately differentiated tumor | Well-moderately differentiated tumor, some high-grade dysplasia, and little stroma, (dirty) necrosis and background | Much well-moderately differentiated tumor epithelium | Little tumor stroma, stroma-low tiles | A couple immune cells, and some (dirty) necrosis, immune-low | Some high-grade dysplasia, some background. A couple vessels with some erythrocytes | Tumor epithelial cells 70%, Stroma 10%, Necrosis 10%, Background 10% |
| 4 | Tumor epithelium | Moderately differentiated tumor - aligned stroma | Well-moderately differentiated tumor, with in between strands of stroma, some immune cells and a few spots of dirty necrosis | Well-moderately differentiated tumor | Some strands of aligned tumor stroma, stroma-low tiles | Some immune cells, and a few spots of dirty necrosis, immune-high overall | N/A | Tumor epithelial cells 60%, Stroma 30%, Immune cells 10% |
| 5 | Necrosis | Necrotic | Mostly (dirty) necrosis, some tumor epithelial cells and some vessels, with background | A couple isolated tumor cells | Some aligned tumor stroma strands, stroma-low tiles | Primarily tiles containing (dirty) necrosis | Some vessels with dirty necrosis and some background or similarly fatty tissue or mucin | Necrosis 80%, Tumor epithelial cells 10%, Background 10% |
| 6 | Healthy and dysplastic colon tissue | Inflamed healthy colon tissue | Healthy and some dysplastic colon tissue, with many immune cells in proliferative and often vascularized stroma, aspect of inflammation | No tumor epithelium | No tumor stroma, only inflamed and vascularized 'healthy' stroma | Many immune cells | Primarily healthy and some low-grade dysplastic colon tissue with vascularized and proliferative aspect of stroma in between, some inflammation | Healthy or dysplastic colon tissue 70%, Immune cells 30% |
| 7 | Necrosis | Avital tumor with necrosis | Tiles are approx. half background, some stroma and necrosis, but mostly moderately differentiated tumor | Mostly moderately differentiated tumor epithelium | Some strands of aligned tumor stroma, stroma-low tiles | Some immune cells and necrosis, immune-high | Much background | Tumor epithelial cells 40%, Background 30%, Stroma 10%, Necrosis 20% |
| 8 | Tumor epithelium | Mixed tumor epithelial - infiltrated stroma | Some variation, mostly tumor and some stroma in different levels of differentiation and organization. | Mostly moderatly differentiated tumor epithelium | Some stroma on mostly stroma-low tiles, some aligned strands but more disorganized in between tumor epithelium | Part immune cells, immune-high | Some background and spots of erythrocytes | Tumor epithelial cells 40%, Stroma 20%, Immune cells 20%, Background 10%, Erythrocytes 10% |
| 9 | Tumor epithelium | Moderately differentiated tumor - aligned stroma | Some background, but mostly moderately differentiated tumor and some stroma in between, a few spots of mucin | Mostly well-moderately differentiated tumor epithelium | Some aligned strands of stroma, tiles are stroma-low | A few immune cells, immune-low | Some background, some mucin | Tumor epithelial cells 70%, Stroma 20%, Background 10% |
| 10 | Muscle tissue | Muscle tissue (longitudinal fibers) | Muscle tissue fiber strands as well as similar looking aligned strands of stroma. Some vessels with erythrocytes, and immune cells in between | No tumor epithelium | Some tiles contain strands of aligned stroma, then the tile is stroma-high | Some immune cells, immune-low in stroma tiles | Mostly muscle tissue, longitudinal fibers and some vessels | Muscle 60%, Stroma 30%, Immune cells 10% |
| 11 | Tumor stroma | Vascularized and infiltrated stroma | Mostly much tumor stroma with neovascularization, and with many immune cells in between, potentially parts of lymphoid tissue. Few isolated tumor epithelial cells | Few isolated tumor epithelial cells | Mostly much and disorganized tumor stroma with neovascularization, stroma-high | High amount of immune cells, potentially part of lymphoid tissue. Immune-high | A couple tiles with a bit background or fatty tissue | Stroma 70%, Immune cells 30% |
| 12 | Mucinous | Mucinous tumor | Some background in otherwise mucinous tumor containing tiles. Notably high contrast in colors, little to no stroma | Predominantly mucinous tumor epithelium | Some stroma, disorganized spots. Stroma-low | Some immune cells, immune-low | Mostly mucus, some background and one tile with fatty tissue | Tumor epithelial cells 40%, Mucin 30%, Background 20%, Stroma 10% |
| 13 | Immune cells | Infiltrated stroma | The mostly aligned tumor stroma is highly infiltrated with immune cells, with some well-moderately differentiated tumor epithelial tissue and some background | Well differentiated tumor epithelium, high-grade dysplasia | Mostly aligned, straight strands of tumor stroma on the stroma-low tiles | High amount of immune cells in stroma, potentially part of lymphoid tissue. Also some dirty necrosis. Immune-high | N/A | Immune cells 30%, Stroma 30%, Tumor epithelial cells 30%, Necrosis 10% |
| 14 | Mucinous | Mucinous tumor | Tumor epithelial cells with some stroma, on tiles consisting approx. half of mucin, some background | Predominantly mucinous tumor epithelium | A part disorganized tumor stroma, some aligned strands. Officially stroma-high tiles, excluding the mucus | A few immune cells, immune-low | Mostly mucus, some background | Mucin 40%, Tumor epithelial cells 30%, Stroma 20%, Background 10% |
| 15 | Fatty tissue | Fatty tissue | Much fatty tissue, some vessels (also round shape), strands of mostly aligned stroma, some immune cells | No tumor epithelium | Strands of aligned stroma, potentially not only tumor-induced but some healthy as well | A part immune cells, some lymphoid tissue, in stroma then immune-high overall | Predominantly fatty tissue on the tiles, and some other round shapes like vessels, some with erythrocytes, and a spot of mucin | Fatty tissue 60%, Stroma 30%, Immune cells 10% |
| 16 | Tumor epithelium | Well-differentiated tumor | Mostly well-moderately differentiated tumor, some stroma in between, and dirty necrosis | Well differentiated tumor epithelium, high-grade dysplasia | Some aligned strands of stroma, tiles are stroma-low | Some immune cells, some dirty necrosis, immune-low | Some background | Tumor epithelial cells 60%, Stroma 20%, Necrosis 10%, Background 10% |

Table 1: Continued

| HPC | Super-cluster | Label | General description | Tumor epithelium | Tumor stroma | Immune cells | Other | HPC tissue type composition |
|---|---|---|---|---|---|---|---|---|
| 17 | Tumor stroma | Infiltrated aligned stroma-high | Some immune cells and tumor epithelial cells in the much most prevalent aligned strands of stroma (stroma-high). Sometimes a neural bundle and a few spots more similar to muscle fibers | Some parts tumor epithelium, well-differentiated | Predominantly tumor stroma, mostly aligned strands on stroma-high tiles | Part immune cells infiltrated in tumor stroma, immune-high overall | Some spots background, a couple tiles with some muscle tissue. A sporadic neural bundle | Stroma 60%, Immune cells 20%, Tumor epithelial cells 10%, Muscle tissue 10% |
| 18 | Immune cells | Infiltrated stroma | Cluster abundant in cells. Much stroma, mostly disorganized, with many immune cells and some isolated tumor cells | Part isolated tumor epithelial cells | Predominantly tumor stroma, mostly disorganized stroma-high tiles | Many immune cells infiltrated in tumor stroma, immune-high | N/A | Stroma 60%, Immune cells 30%, Tumor epithelial cells 10% |
| 19 | Tumor stroma | Edematous disorganized stroma | Much edematous, mostly disorganized stroma, almost no immune cells but with tumor epithelium, some necrosis | Some parts tumor epithelium, well-differentiated | Much edematous tumor stroma, mostly disorganized stroma-high tiles | Few immune cells, some (dirty) necrosis, immune-low | A few spots mucin | Stroma 70%, Tumor epithelial cells 30% |
| 20 | Tumor epithelium | Disorganized and vascularized stroma-low | Some background but mostly moderately differentiated tumor epithelial cells in disorganized stroma, some immune cells | Well-moderately differentiated tumor | Mostly disorganized stroma, stroma-low tiles, some vascularization | A part immune cells, some (dirty) necrosis, immune-low mostly | Some background and a few spots mucin, a slight signet cell component | Tumor epithelial cells 50%, Stroma 30%, Immune cells 10%, Background 10% |
| 21 | Tumor stroma | Vascularized stroma | Aligned strands of much loose stroma with neovascularization (or small white shapes similar to vessels), sometimes some muscle fibers, with immune cells and isolated tumor cells in between | A couple isolated tumor cells | Predominantly aligned tumor stroma strands, stroma-high, vascularized (or other round shapes) | Part immune cells infiltrated in tumor stroma, some parts immune-high | Some background, a few spots of fatty tissue. A couple tiles muscle tissue in stead of stroma | Stroma 70%, Immune cells 10%, Tumor epithelial cells 10%, Muscle tissue 10% |
| 22 | Immune cells | Infiltrated stroma | Many immune cells in tumor-induced and healthy stroma, with some tumor or healthy colon tissue and some background | Some well differentiated tumor epithelium, high-grade dysplasia | Healthy and tumor stroma, which is stroma-high | High number of immune cells, some necrosis, immune-high | Some healthy and dysplastic tissue | Immune cells 30%, Stroma 30%, Tumor epithelial cells 20%, Background 10%, Healthy and dysplastic colon tissue 10% |
| 23 | Healthy and dysplastic colon tissue | Dysplastic colon tissue | Mostly dysplastic colon tissue (adenoma), with many immune cells in between and much background and mucin | A couple isolated tumor cells | Dysplastic stroma | Many immune cells in dysplastic stroma | Predominantly dysplastic colon tissue, some background and mucin | Healthy and dysplastic colon tissue 40%, Immune cells 20%, Mucin 20%, Background 20% |
| 24 | Muscle tissue | Vessel-like stroma/ muscle | Approx. half of the tiles contain background, some mucin, the other half is stroma and/or muscle tissue of vessels; large, white round shapes. There is some dirty necrosis | A couple isolated tumor cells | Stroma in the shape of vessels, these are mostly aligned tumor stroma and stroma-high | A few immune cells, some dirty necrosis, immune-low | Muscle tissue, mostly the walls from vessels and background, a spot of erythrocytes within these vessels, some mucin | Muscle tissue 30%, Stroma 30%, Background 30%, Mucin 10% |
| 25 | Necrosis | Moderately differentiated tumor - necrotic | Moderately differentiated tumor epithelial with (dirty) necrosis, and some stroma in between. Some background and mucin as well | Mostly well differentiated tumor epithelial cells | Some strands of aligned tumor stroma, stroma-low tiles | Part immune cells in (dirty) necrosis, otherwise immune-low | In general busy and disorganized tiles, round shapes, some background, some mucin | Tumor epithelial cells 50%, Necrosis 20%, Stroma 10%, Mucin 10%, Background 10% |
| 26 | Tumor epithelium | Infiltrated poor-undifferentiated tumor | Cluster rich in cells. Tiles with solid growing poor-undifferentiated tumor, others immune cells and some stroma in between. Potentially microsatellite instability | Tiles with solid growing poor-undifferentiated tumor epithelium | A couple strands of aligned tumor stroma, stroma-low tiles | Many immune cells, some tiles only containing immune cells, immune-high | Cluster rich in cells, one tile a slight signet cell component, a spot background | Tumor epithelial cells 80%, Immune cells 10%, Stroma 10% |
| 27 | Tumor epithelium | Moderately differentiated tumor - aligned stroma | Some background or mucin. Well-moderately differentiated tumor epithelial cells in some aligned strands of stroma and some immune cells | Well differentiated tumor epithelium, high-grade dysplasia | Some strands of aligned tumor stroma, stroma-low tiles | Some immune cells, also in (dirty) necrosis, overall immune-high | Some background | Tumor epithelial cells 50%, Stroma 30%, Immune cells 10%, Background 10% |
| 28 | Muscle tissue | Muscle (longitudinal fibers) | Mostly muscle tissue, longitudinal sections, sometimes aligned stroma strands and some background. A few tumor epithelial cells and neural bundles | A couple isolated tumor cells | A couple tiles contain strands of aligned stroma, these are stroma-high | A few immune cells, immune-low in stroma tiles | Predominantly muscle tissue in longitudinal fibers, some background or a spot of fatty tissue. A sporadic neural bundle | Muscle tissue 70%, Stroma 20%, Background 10% |
| 29 | Muscle tissue | Muscle (longitudinal fibers) | Mostly longitudinal sectioned muscle tissue and/or stroma strands, a bit edematous, some vessels, a neural bundle, not many immune cells | A couple isolated tumor cells | A couple tiles contain strands of aligned stroma, then the tile is stroma-high | A few immune cells, immune-low in stroma tiles | Predominantly muscle tissue in longitudinal fibers, some background or a spot of fatty tissue. A sporadic neural bundle. Small vessels often in between | Muscle tissue 80%, Stroma 20% |
| 30 | Immune cells | Mixed, immune-high | Aligned stroma strands and well-moderately differentiated tumor epithelial cells, immune cells in between, some dirty necrosis | Well differentiated tumor epithelium, high-grade dysplasia | Aligned stroma strands on mostly stroma-low tiles | Many immune cells infiltrated in tumor stroma, some spots of (dirty) necrosis | N/A | Tumor epithelial cells 30%, Stroma 30%, Immune cells 30%, Necrosis 10% |
| 31 | Immune cells | Mixed, infiltrated stroma | Often aligned stroma with many immune cells and moderately differentiated tumor epithelium. Often some background, some necrosis | Well differentiated tumor epithelium, high-grade dysplasia | Aligned, loose stroma strands on mostly stroma-low tiles | Part immune cells often infiltrated in tumor stroma, immune-high | Some background, a couple spots of mucin or fatty tissue | Stroma 40%, Immune cells 20%, Tumor epithelial cells 20%, Background 20% |

Table 1: Continued

| HPC | Super-cluster | Label | General description | Tumor epithelium | Tumor stroma | Immune cells | Other | HPC tissue type composition |
|---|---|---|---|---|---|---|---|---|
| 32 | Muscle tissue | Muscle (axial section) | Axial sections of muscle tissue, some disorganized strands of stroma and some background. Also, some tiles that contain erythrocytes, similar looking to the muscle fiber sections | No tumor epithelium | No tumor stroma | A few immune cells | Tiles contain mostly muscle tissue, with the fibers in axial sections. Some are part of a vessel wall, with sometimes also erythrocytes in vessels. Some background | Muscle tissue 70%, Erythrocytes 10%, Stroma 10%, Background 10% |
| 33 | Muscle tissue | Muscle (longitudinal fibers) | Muscle fibers in longitudinal sections or aligned stroma strands, with much background and fatty tissue, some vessels and only few immune cells | A couple isolated tumor cells | Some tiles contain strands of aligned stroma, then the tile is stroma-high | A few immune cells, immune-low in stroma tiles | Some background, some fatty tissue, in between loose bundles of longitudinal muscle fibers, some muscle tissue as part of vessels | Muscle tissue 40%, Stroma 30%, Background 20%, Fatty tissue 10% |
| 34 | Muscle tissue | Muscle (axial section) | Mostly muscle tissue in axial plane, some tiles disorganized tumor stroma, with some vessels; overall few nuclei or immune cells | A couple isolated tumor cells | A couple tiles contain strands of disorganized tumor stroma, these are stroma-high | A few immune cells, immune-low in stroma tiles | Predominantly axial sections of muscle tissue, a few spots of background or mucus | Muscle tissue 70%, Stroma 30% |
| 35 | Immune cells | Immune cells | Mostly immune cells in stroma or lymphoid tissue, with some tumor epithelial cells or necrosis | A couple isolated tumor cells | Tumor- and healthy stroma, stroma-high tiles | Highly infiltrated (tumor) stroma or necrosis, sometimes part of lymphoid tissue, immune-high | Some background, a few spots of mucin or fatty tissue, a few spots of dysplastic colon tissue | Immune cells 60%, Stroma 20%, Tumor epithelial cells 10%, Background 10% |
| 36 | Necrosis | Avital tumor with necrosis | Loose and pieces of tissue, mostly avital tumor epithelium or necrosis, in tiles with much background, some strands of stroma | Loose and pieces of tissue, mostly avital tumor epithelium, well-moderately differentiated | Some spots of stroma, mostly disorganized, stroma-low tiles | Immune cells in much present necrosis | Much background, a few spots of mucin | Necrosis 30%, Tumor epithelial cells 30%, Background 30%, Stroma 10% |
| 37 | Healthy and dysplastic colon tissue | High-grade dysplasia - adenoma | Adenoma or well differentiated tumor, with much background, some mucin, and few spots of dirty necrosis | Adenoma, high-grade dysplasia or well differentiated tumor epithelium | A few strands of aligned (tumor) stroma, stroma-low | Some immune cells in dysplastic and tumor stroma, a spot of dirty necrosis | Some background and/or mucin | Healthy or dysplastic colon tissue 60%, Stroma 10%, Immune cells 10%, Background 10%, Mucin 10% |
| 38 | Mucinous | Mucinous tumor stroma | Mucin, mostly from tumors with disorganized tumor stroma, some background | A couple isolated tumor cells | Some disorganized strands of tumor stroma in between the mucin, tiles are stroma-high | A few immune cells, immune-low | Predominantly mucin | Mucin 80%, Stroma 20% |
| 39 | Healthy and dysplastic colon tissue | Healthy colon tissue | Mostly clear, healthy colon tissue with mucin in goblet cells and the villi, a small component dysplasia, some background and immune cells in between | No tumor epithelium | No tumor stroma | Some immune cells in healthy stroma, normal | Predominantly healthy colon tissue, some background and/or mucin | Healthy or dysplastic colon tissue 80%, Background 10%, Mucin 10% |
| 40 | Tumor stroma | Disorganized stroma, growing infiltratively | Polymorphic and poorly differentiated tumors in much prevalent edematous and disorganized stroma, growing infiltratively, a few immune cells | Part polymorphic and poorly differentiated tumor epithelium, growing infiltratively | Much prevalent edematous and disorganized tumor stroma, often vascularized, stroma-high | Part immune cells, immune-low | N/A | Stroma 60%, Tumor epithelial cells 30%, Immune cells 10% |
| 41 | Tumor stroma | Disorganized and loose stroma | Heterogeneous cluster. Some background and/or (loose) stromal tissue, not all tumor-associated, with some vessels and immune cells, a few spots of mucin | A couple isolated tumor cells | Predominantly loose aligned stroma fibers, some tumor-induced, then stroma-high. Vascularized | Part immune cells, immune-low | Some tiles contain muscle tissue from vessels, some background, a few spots of mucin | Stroma 70%, Muscle tissue 10%, Background 10%, Immune cells 10% |
| 42 | Tumor epithelium | Moderately differentiated tumor | Mostly moderately differentiated tumor epithelial with a few aligned strands of stroma in between. Some background and a few artefacts | Mostly well-moderately differentiated tumor epithelium | Some aligned strands of stroma, tiles are stroma-low | Few immune cells, some spots (dirty) necrosis, immune-low | Some background | Tumor epithelial cells 70%, Stroma 20%, Background 10% |
| 43 | Necrosis | Necrotic | Necrosis with anucleic cells, or fields with mostly immune cells. Much background and some tumor epithelial cells | Some moderately tumor epithelium, some isolated tumor epithelial cells | A few strands of aligned (tumor) stroma | Immune cells in much present necrosis | Some background | Necrosis 50%, Immune cells 20%, Background 20%, Tumor epithelial cells 10% |
| 44 | Tumor stroma | Disorganized, immune-low stroma (desert-type) | Mostly aligned stroma, desert type with few nuclei and immune cells, potentially some not tumor-induced with a few isolated tumor epithelial cells | A couple isolated tumor cells | Much prevalent edematous and disorganized tumor stroma, often vascularized, stroma-high | A few immune cells, immune-low | A couple tiles contain longitudinal muscle fibers | Stroma 90%, Immune cells 10% |
| 45 | Tumor epithelium | Poor-undifferen-tiated tumor | Poor-undifferentiated tumor epithelium, with some disorganized stroma, some background | Much poor-undifferentiated tumor epithelium | Some disorganized tumor stroma, stroma-low | A few immune cells, immune-low | Some background | Tumor epithelial cells 70%, Stroma 20%, Background 10% |
| 46 | Tumor epithelium | Infiltrated undifferen-tiated tumor | Cluster rich in cells. Solid growing, undifferentiated tumor, with some immune cells. Potentially micro-satellite instability. Component signet cell carcinoma and squamous cells | Much poor-undifferentiated tumor epithelium | A few strands of aligned (tumor) stroma, stroma-low | Many immune cells, also a couple tiles with only immune cells, immune-high | A tile with slight component signet cell, one with necrosis | Tumor epithelial cells 70%, Immune cells 20%, Stroma 10% |

**a**

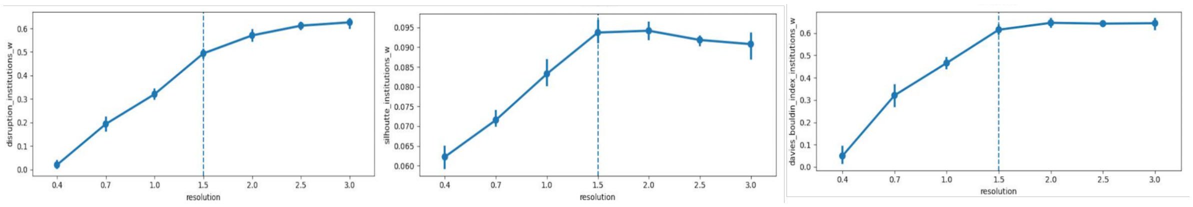


Leiden resolution optimization using unsupervised methods

**b**

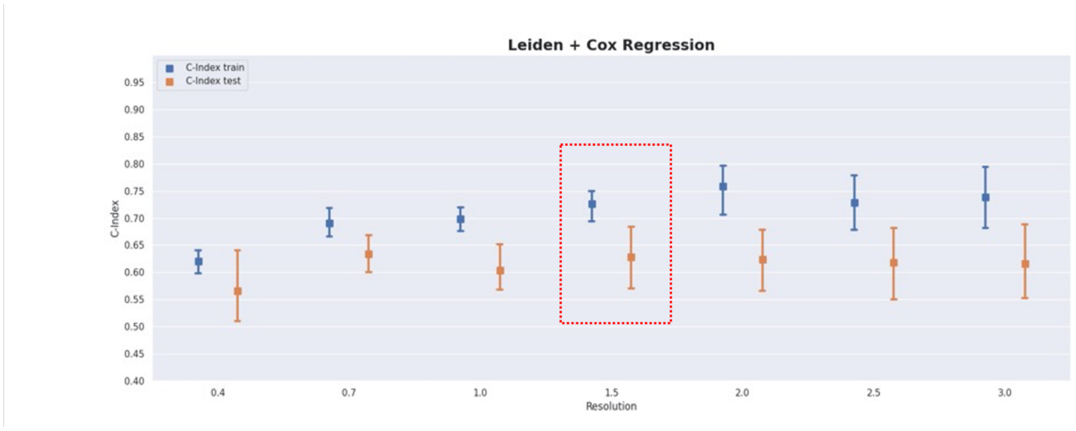


Leiden resolution optimization on OS prediction from regularized Cox regression in TCGA

Figure 1: **Optimization of Leiden resolution** (a) Disruption score, Silhouette score, and Daves-Boundin index weighted by mean percentage of the institution presence in each HPC reached a consensus on optimal Leiden resolution at 1.5. (b) We trained regularized Cox regressions including all 47 HPCs for each Leiden resolution using 5-fold CV in TCGA. The Leiden resolution 0.7 and 1.5 showed the highest validation c-index. We chose Leiden 1.5 as the optimal resolution because it led to more explorable HPCs. CV, cross-validation. HPC, histomorphological phenotype cluster. OS, overall survival. TCGA, The Cancer Genome Atlas.

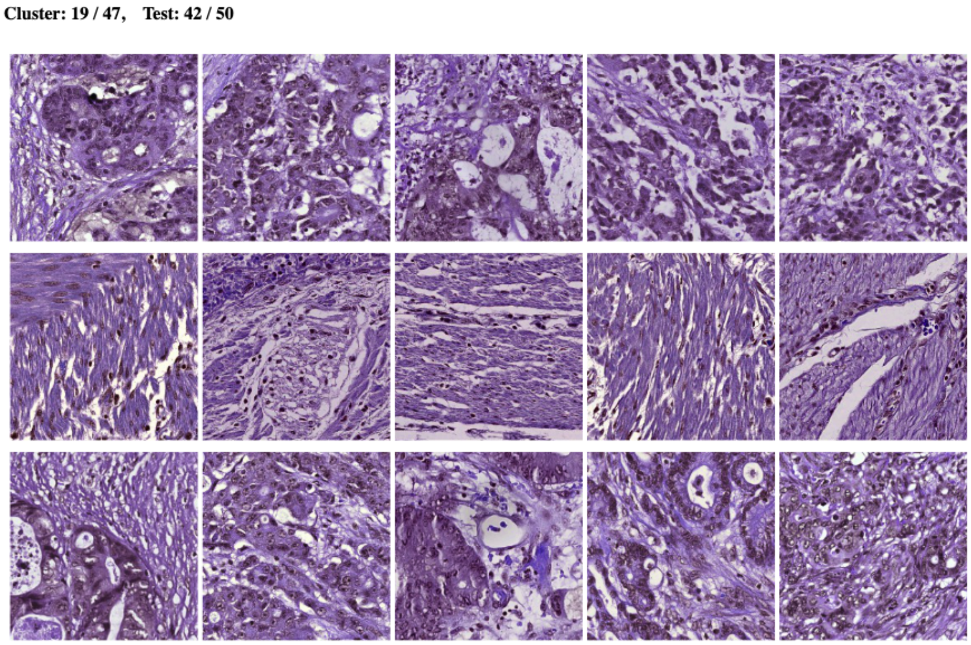


Figure 2: **Objective test for all HPCs.** Screenshot of the online objective test. Three rows of tiles were presented at each test, with 2 rows were from the same HPC and the other row from a randomly selected different HPC.

**a HPC-based classifier in five-fold CV in TCGA**

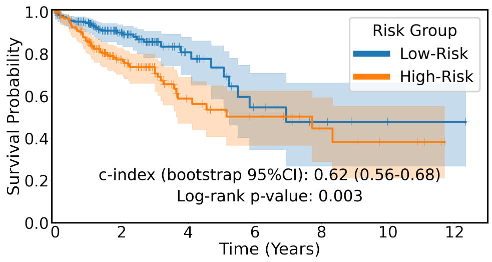


**b HPC-based classifier in the external AVANT control group**

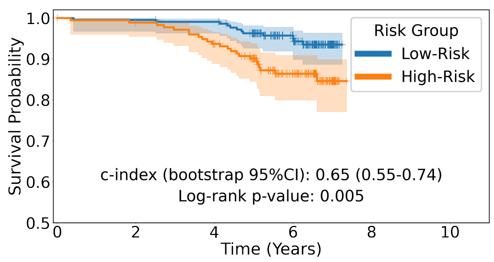


**c Clinical baseline classifier in the external AVANT control group**

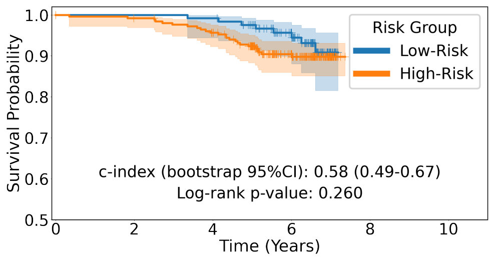


**d HPC-based classifier in five-fold CV in AVANT experimental group**

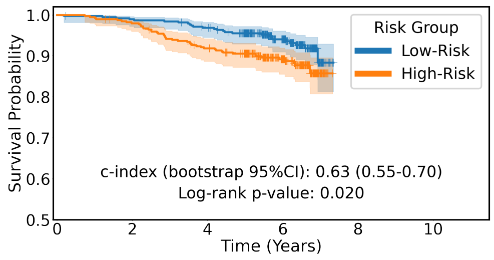


Figure 3: **Regularized Cox regression for OS prediction.** (a) Kaplan-Meier plot showing the HPC-based classifier stratifying OS risk in the five-fold CV in TCGA (log-rank P=0.003). (b) The HPC-based classifier stratifying OS risk in the external AVANT control test set (log-rank P=0.005). (c) The clinical baseline model (predictors including age, sex, tumor-stroma ratio, AJCC TNM staging) stratifying OS risk in the independent AVANT control test set (log-rank P=0.260). (d) The HPC-based classifier stratified OS risk AVANT-experimental group (log-rank P=0.020). CV, cross-validation. HPC, histomorphological phenotype cluster. OS, overall survival. TCGA, The Cancer Genome Atlas.
